# An analytical approach to bistable biological circuit discrimination using real algebraic geometry

**DOI:** 10.1101/008581

**Authors:** Dan Siegal-Gaskins, Elisa Franco, Tiffany Zhou, Richard M. Murray

## Abstract

Biomolecular circuits with two distinct and stable steady states have been identified as essential components in a wide range of biological networks, with a variety of mechanisms and topologies giving rise to their important bistable property. Understanding the differences between circuit implementations is an important question, particularly for the synthetic biologist faced with determining which bistable circuit design out of many is best for their specific application. In this work we explore the applicability of Sturm’s theorem—a tool from 19th-century real algebraic geometry—to comparing “functionally equivalent” bistable circuits without the need for numerical simulation. We first consider two genetic toggle variants and two different positive feedback circuits, and show how specific topological properties present in each type of circuit can serve to increase the size of the regions of parameter space in which they function as switches. We then demonstrate that a single competitive monomeric activator added to a purely-monomeric (and otherwise monostable) mutual repressor circuit is sufficient for bistability. Finally, we compare our approach with the Routh-Hurwitz method and derive consistent, yet more powerful, parametric conditions. The predictive power and ease of use of Sturm’s theorem demonstrated in this work suggests that algebraic geometric techniques may be underutilized in biomolecular circuit analysis.

## 1 Introduction

The field of synthetic biology has rapidly matured to the point where it is now possible to produce complex synthetic networks with prescribed functions and level of performance [1]. As in other fields of engineering, advances have been enabled by the use of small interchangeable modules that are “functionally equivalent” from an input-output perspective [2]. Bistable circuits—which play a role in essential biological processes including cell fate specification [3], cell cycle progression [4], and apoptosis [5]—make up a particularly large and diverse functionally equivalent set [6]. Effectively characterizing and comparing these biocircuits is crucial for determining which design is in some sense optimal for a particular context.

Ordinary differential equation (ODE) models can be powerful tools for identifying and contrasting biocircuits’ “dynamic phenotypes” (see, e.g., [7]). Numerical simulations are often used in the analysis of these models; however, analytical criteria that focus on topology can provide a more exact assessment of a circuit’s properties [8, 9]. (Here we use *topology* to mean the particular set of interactions between regulatory parts.) A novel analytical tool that can provide topology-based insights can be found in Sturm’s theorem [10], developed in 1835 as a solution to the problem of finding the number of real roots of an algebraic equation with real coefficients over a given interval. Despite its predictive power, this “gem of 19th century algebra and one of the greatest discoveries in the theory of polynomials” [11] remains an unexploited tool for analysis of biological circuit models.

In this work we demonstrate an approach to bistable circuit discrimination based on Sturm’s theorem that can give the boundaries of the regions of bistability as exact analytic expressions, eliminating the need for numerical simulation. We compare the regions of bistability for two variants of the classic double-negative toggle switch as well as two positive feedback circuits, one of which is based on the bacteriophage λ promoter P__RM__. We then show that while a purely monomeric version of the genetic toggle cannot be bistable, a single competitive activating species added to the circuit leads to bistability in a noncontiguous region of parameter space. Lastly, we use a model of an RNA aptamer-based bistable switch to compare our Sturm’s theorem approach to another based on the control theoretic Routh-Hurwitz method. Overall our results highlight a new use for Sturm’s theorem for identifying potential differences between functionally equivalent bistable biocircuits, and serve as a (re-)introduction to the method as a general tool for studying the kinds of polynomial expressions that often arise when modeling biological systems.

## 2 Mathematical preliminaries

Sturm’s theorem gives the number of distinct real roots of a univariate polynomial *f*(*x*) in a particular interval. To apply the theorem, we must first construct the *Sturm sequence*, a set of polynomials ℱ = {*f*_0_, *f*_1_, …, *f*_*m*_} defined as:

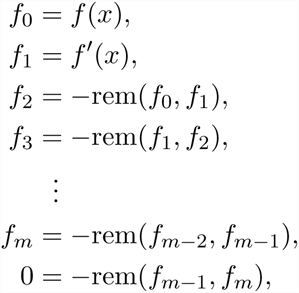

where rem(*f*_*i*_, *f*_*i*+1_) is the remainder of the polynomial long division of *f*_*i*_ by *f*_*i*+1_. The sequence ends at *f*_*m*_, when *f*_*m−1*_ divided by *f*_*m*_ gives a remainder of zero. For a polynomial of degree *n*, there are *m* ≤ *n* + 1 Sturm polynomials in the sequence.

**Theorem 1 (Sturm’s theorem)** *Let f* (*x*) *be a real-valued univariate polynomial and a, b* ∈ ℝ⋃{−∞, +∞}, *with a* < *b and f*(*a*), *f*(*b*) ≠ 0. *Then the number of zeroes of f*(*x*) *in the interval* (*a, b*) *is the difference*

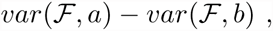

*where* ℱ *is the Sturm sequence of f*(*x*), *and the **variations** var*(ℱ, *a*) *and var*(ℱ, *b*) *are the number of times that consecutive nonzero elements of the Sturm sequence—evaluated at a and b, respectively—have opposite signs.* (Adapted from [12].)

Our approach involves identifying regions of bistability by finding conditions that lead to three steady states, without requiring numerical determination of the exact values or stability of the equilibrium points. While it is in general not possible to draw conclusions on the stability properties of equilibria by simply counting their number, the circuits under consideration enjoy important properties—namely, they are dissipative, their linearizations are positive, and the characteristic polynomials of their Jacobians have all nonconstant terms positive—that allow us to relate their degree and number of equilibria to the stability properties of each equilibrium (see Supporting Information). For such circuits, when three equilibria are present, two of them must be stable and one must be unstable.

## 3 Overview of the method

To determine the region in parameter space in which a particular circuit exhibits bistability, we first find the polynomial f (x) that describes the equilibrium state of the system (where x represents the concentration of a circuit species of interest) and construct its Sturm sequence ℱ = {*f*_0_, *f*_1_, …, *f*_*m*_} as described above. Since we are interested in all possible positive concentrations, *x* → 0 and *x* → +∞ are chosen as the limits of the interval on which Sturm’s theorem is applied. For each limit we generate different sets of polynomial inequalities by independently setting each element of ℱ to be either greater than or less than zero; however, only some of these inequality sets may be combined in such a way to yield a variation difference *var*(ℱ, 0)−*var*(ℱ, +∞) = 3 and thus three steady states. (For example, with *var*(ℱ, 0) = 4, only those complementary sets with var(ℱ, +∞) = 1 need to be considered.) These combinations of inequalities are enumerated and then tested for *logical consistency*, i.e., whether all inequalities can be simultaneously satisfied. It is important to note that, since Sturm’s theorem is only concerned with consecutive nonzero elements and zeroes are ignored, a particular Sturm polynomial may be equal to zero without affecting the total number of sign changes. In cases where zeroes are valid options in the sequence, strict inequalities are made nonstrict (e.g., ‘>’ → ‘≥’).

Symbolic manipulation is done using Mathematica; a sample Mathematica notebook in which the method is applied is included as supplementary material.

## 4 Results

### 4.1 Genetic toggle circuits

A recent study identified a set of eleven *minimal bistable networks* (MBNs), simple two-gene circuits with the capacity for bistability that do not also contain a smaller bistable subcircuit [13]. One of these MBNs, a double-negative toggle switch consisting of two dimeric repressors (Fig. 1A, top), was among the very first synthetic biocircuits built and modeled [14]. This *dimer-dimer* (DD) toggle design has since gone on to be used in a wide range of synthetic biological applications, including the manipulation of fluxes of the *E. coli* metabolic network [15] and as part of a circuit involved in programmed autonomous cellular diversification [16]. A second MBN of particular interest, herein referred to as the *monomer-dimer* (MD) toggle, is a double-negative switch variant in which one of the repressors functions as a monomer (Fig. 1A, bottom). While to our knowledge no MD toggle circuit has been constructed, the components necessary for its implementation exist in the form of monomeric single-chain transcriptional repressor proteins [17], as well as in transcription-activator-like effectors (TALEs) and CRISPR/Cas nucleases that have been engineered as repressors [18,19]. Other exotic toggle-like circuit topologies have also been proposed and/or built (see, e.g., [13, 18, 20–22]). An understanding of the differences in how these various toggles perform can be beneficial, in particular for those circuits that have yet to be built and for which *a priori* knowledge of their behavior could aid in their development.

**Figure 1:**
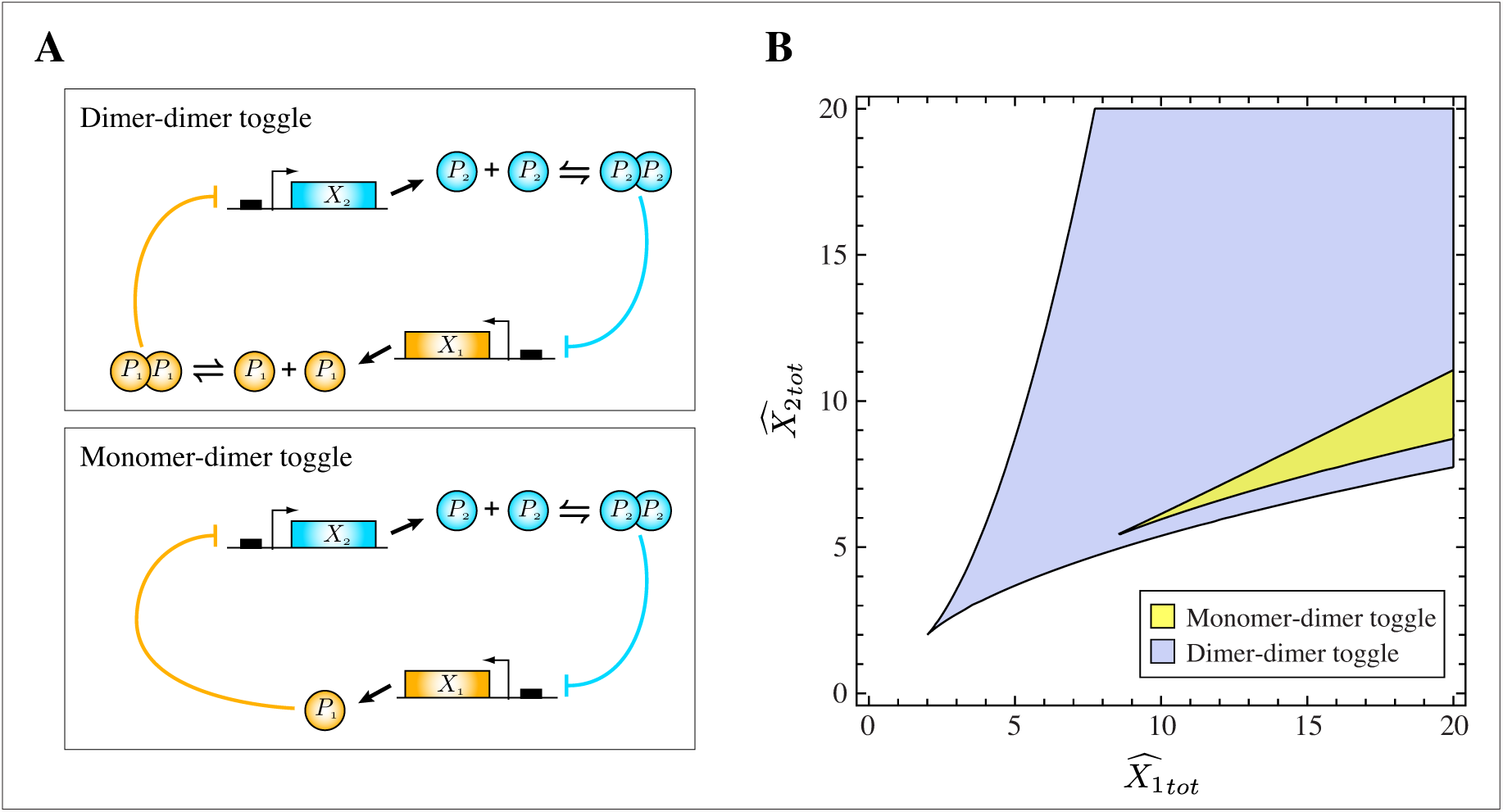
(A) Dimer-dimer (top) and monomer-dimer (bottom) toggle switches. (B) Bistable regions for the monomer-dimer and dimer-dimer toggles. The dimer-dimer toggle exhibits bistability over a much larger range of 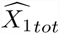 and 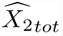.

As a first demonstration of our approach, we apply Sturm’s theorem to the DD and MD toggle circuits. Beginning with a chemical reaction network formulation and assuming mass-action kinetics we derive ODE models of the two toggles (Eqs. (S1) and (S2)). At equilibrium the concentrations of *P*_1_ and *P*_2_ in the MD system are given by

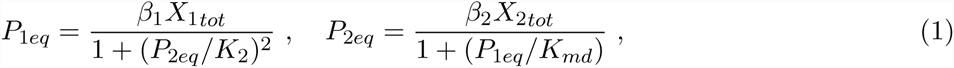

and in the DD system,

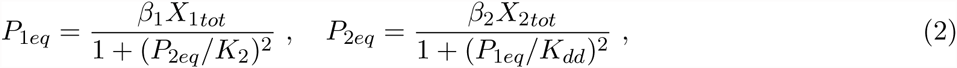

where *X*_*itot*_ is the total concentration of gene *i*, *β_i_* = *k*_*basi*_/*k*_*degi*_ is the ratio of basal production and degradation rate constants for protein *i*, *K*_*xd*_ = {*K*_*md*_, *K*_*dd*_} is the Michaelis constant for *P*_1_, and *K*_2_ is the Michaelis constant for *P*_2_. (Recall that Michaelis constants represent the repressor concentrations that yield 50% of the maximum production rate of their target proteins and are determined by the protein-DNA and dimerization dissociation constants; see Supporting Information for details.) Systems (1) and (2) may be written in terms of *P*_1*eq*_ alone, as:

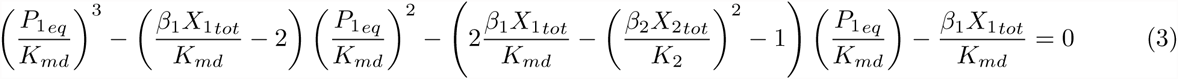

and

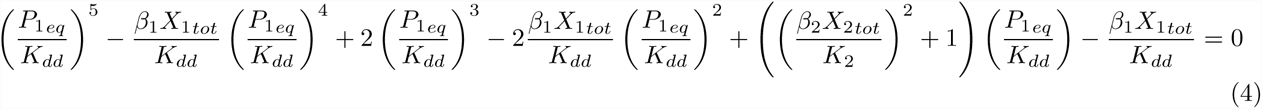

With the following scaling of the DNA and protein concentrations:

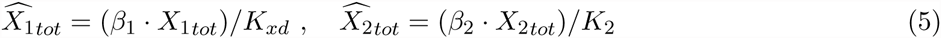

and

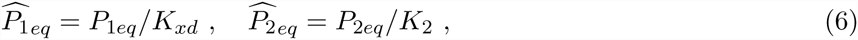

we may write Eqs. (3) and (4) as non-dimensional polynomials in 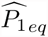:

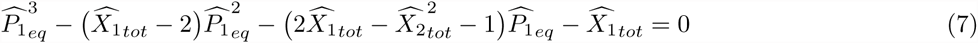

and

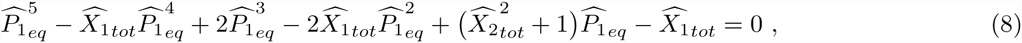

for the MD and DD toggles, respectively. Every positive root of these equilibrium polynomials gives a positive steady state concentration for every other circuit component as well.

We may now determine the regions of bistability in the plane of total (non-dimensionalized) DNA concentrations 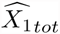 and 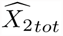. As outlined in the ‘Overview’ section, the first step is the construction of a Sturm sequence *F* for each equilibrium polynomial. These sequences are given in the Supporting Information. It can be seen that, for the DD toggle, some ofvthe polynomials in the sequence contain a 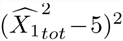 term that renders them indeterminate when 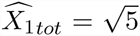. (The same term is responsible for one of the polynomials going to zero, thus terminating the sequence prematurely.) To avoid these problems it is necessary to generate two different sequences for the DD toggle: one for which we assume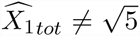 in Eq. (8), and another with 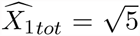. In this latter case the DD toggle equilibrium polynomial Eq. (8) becomes

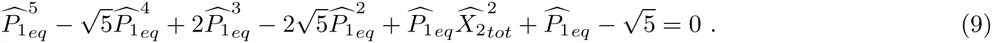

Following construction of the toggle Sturm sequences, we evaluate each at 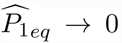 and 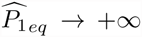 and find the conditions that give a variation difference of 3. The MD toggle Sturm sequence ℱ_*md*_ has a maximum possible variation of 3 and only one combination of inequalities that can give rise to bistability: when *var*(ℱ_*md*_, 0) = 3 and *var*(_*md*_, +) = 0. In contrast, the DD toggle sequence ℱ_*dd*_ could in principle yield as many as five positive steady states; however, only three are admitted as there are no combinations of inequalities that have a variation difference of 5 or 4 and are logically consistent. (See ‘Overview’ section for details.) Inequality sets for the DD toggle are listed (in the compact form {± ± … ±}) in Tables 1 and 2.

**Table 1:**
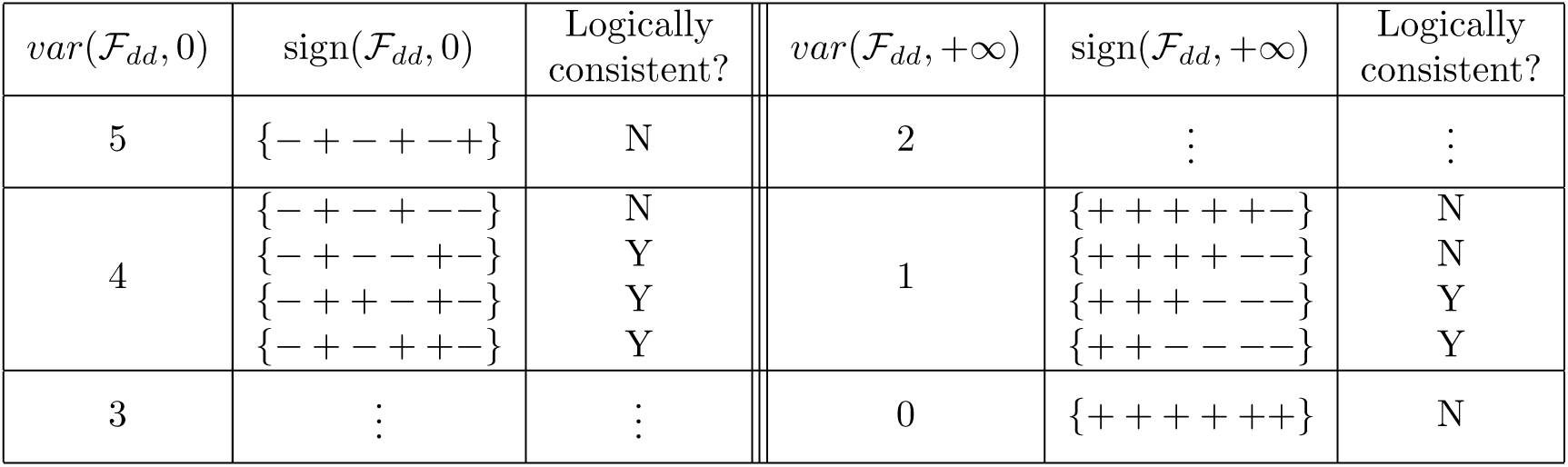
Sturm sequence inequality sets for the DD toggle when 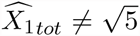. The signs of the first two polynomials are fixed at 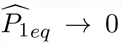 and 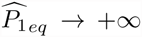. Neither var(ℱ_*dd*_, 0) = 5 nor var(ℱ_*dd*_, +∞) = 0 represent logically consistent sets.

| $var(\mathcal{F}_{dd}, 0)$ | $sign(\mathcal{F}_{dd}, 0)$ | Logically consistent? | $var(\mathcal{F}_{dd}, +\infty)$ | $sign(\mathcal{F}_{dd}, +\infty)$ | Logically consistent? |
| --- | --- | --- | --- | --- | --- |
| 5 | $\{- + - + - +\}$ | N | 2 | ⋮ | ⋮ |
| 4 | $\{- + - + --\}$ | N | 1 | $\{+ + + + + -\}$ | N |
| | $\{- + - - + -\}$ | Y | | $\{+ + + + - -\}$ | N |
| | $\{- + + - + -\}$ | Y | | $\{+ + + - - -\}$ | Y |
| | $\{- + - + + -\}$ | Y | | $\{+ + - - - -\}$ | Y |
| 3 | ⋮ | ⋮ | 0 | $\{+ + + + + +\}$ | N |

**Table 2:** Sturm sequence inequality sets for the DD toggle when 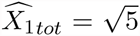. The signs of the first two and three polynomials are fixed at 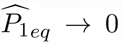 and 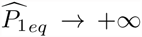, respectively. The set with var(F_*dd*_, 0) = 4 is not logically consistent.

| $var(\mathcal{F}_{dd}, 0)$ | $sign(\mathcal{F}_{dd}, 0)$ | Logically<br>consistent? | $var(\mathcal{F}_{dd}, +\infty)$ | $sign(\mathcal{F}_{dd}, +\infty)$ | Logically<br>consistent? |
| --- | --- | --- | --- | --- | --- |
| 4 | $\{- + - + -\}$ | N | 1 | ⋮ | ⋮ |
| 3 | $\{- + - + +\}$ | N | 0 | $\{+ + + + +\}$ | Y |
| | $\{- + + - +\}$ | Y | | | |
| | $\{- + - - +\}$ | N | | | |

The valid inequality sets were reduced with Mathematica to yield analytic expressions for the regions of bistability in the plane of 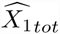 and 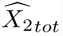. For the MD toggle, this bistability region is found to be

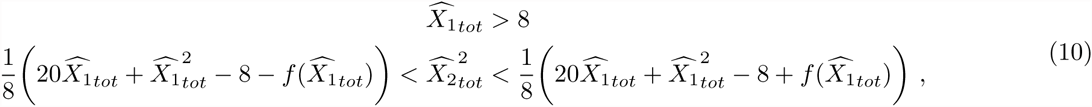

with 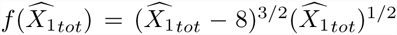. The DD toggle inequality sets combine to give a continuous region of bistability that is most easily written as the intersection of

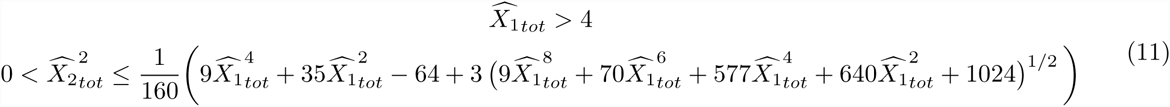

and

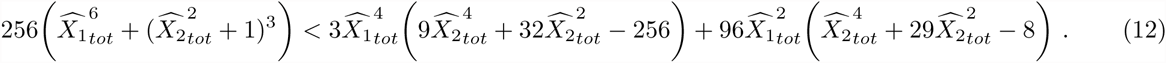

It is important to note that, with the exception of the DD toggle at 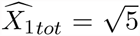 (which was treated by analyzing a second Sturm sequence, as described above), none of the potential zeroes in the Sturm polynomial denominators required special treatment nor did they present any problems in determining the regions of bistability.

Our analysis shows that the DD toggle operates as a bistable switch over a substantially greater range of (non-dimensionalized) DNA concentrations than does the MD toggle (Fig. 1B), indicating that the DD topology is more functionally robust to variations in DNA concentrations and rate parameters. Furthermore, the DD switch can operate with significantly lower values of 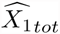 and 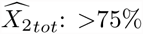 and >50% less, respectively. Worth highlighting is the fact that, through application of our method to the non-dimensionalized versions of the original systems, we have demonstrated that the *function* of bistability is determined by a particular combination of the system parameters— gene concentrations, protein basal production and degradation rate constants, and Michaelis constants—and thus there is no single parameter more important than any other for achieving bistability.

#### Computational support

The approach presented here does not require any numerical simulation; however, some computational validation of our results may be considered of value. For both toggle circuits, and for each of the valid combinations of sign(ℱ, 0) and sign(ℱ, +∞), 1000 random values of 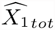 and 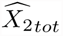 were selected from inside and outside of the predicted bistable regions and plugged in to the appropriate equilibrium polynomial (Eq. (7) or (8)) that were then solved numerically. In all cases the number of equilibria found matched the number determined by Sturm’s theorem: three equilibria were found inside the bistable regions and only one equilibrium was found outside.

We may also check the stability of the various steady states using the circuits’ Jacobian J and characteristic polynomial *p*_*J*_ (λ) = det (λ*I* − *J*). It was recently shown that if all off-diagonal components of the Jacobian are nonnegative (i.e., it is a Metzler matrix), or if the Jacobian may be transformed to have such a form, then any equilibrium is unstable if and only if the constant term of *p*_*J*_ (λ) has a sign opposite to that of all other terms in *p*_*J*_ (λ) [20]. We use this condition on the constant term of *p*_*J*_ (λ) to confirm that each bistable solution set contains one and only one unstable steady state.

The inequalities that satisfy the *p*_*J*_ (λ) constant term condition are:

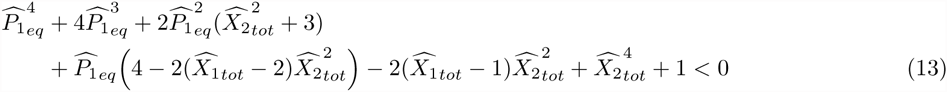

and

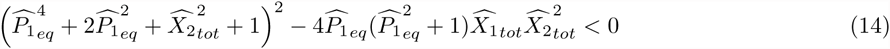

for the MD and DD toggles, respectively. For each bistable solution set found we substituted the values 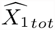, 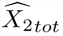 and 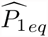, into Eqs. (13) and (14) and confirmed that only one of the three solutions satisfies the appropriate instability condition.

It is worth noting that the time required to test 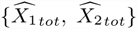 pairs scales linearly with the number of pairs, so while testing a small number can be done relatively quickly, as the number of pairs becomes appreciable the time can be significant—up to 3 hours to test 600,000 random values of 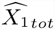 and 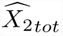.

## 4.2 Bistable single-gene circuits

The single gene system consisting of bacteriophage λ repressor and its promoter P_RM_ (with its three operator sites OR1, OR2, and OR3) also exhibits bistability [23]. Using the dimensionless model given in [23] we have that the steady state concentration of protein satisfies

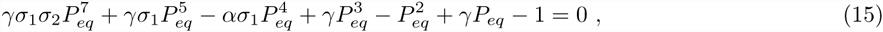

where *γ* is the rescaled degradation rate constant, α represents the increase in protein production resulting from dimer binding to OR2, and σ_1_ and σ_2_ are the relative (to OR1) affinities for OR2 and the negatively-regulating OR3, respectively. (For simplicity we set the gene copy number equal to one.) With σ_1_ = 2 and σ_2_ = 0.08 [23], the associated Sturm sequence F_P_RM has only two inequality sets with var(ℱ_P_RM__, 0) = 5 and one set with var(F_P_RM__, +∞) = 2 that are logically consistent and together give bistability (Table 3).

**Table 3:** Sturm sequence inequality sets for the λ repressor–P_RM_ system. The first four Sturm polynomials have fixed signs at *P*_*eq*_ = 0 and *P*_*eq*_ = +∞. Neither *var*(ℱ_P_RM__, 0) = 6 nor *var*(ℱ_P_RM__, +∞) = 1 represent logically consistent sets.

| $var(\mathcal{F}_{P_{RM}}, 0)$ | $sign(\mathcal{F}_{P_{RM}}, 0)$ | Logically consistent? | $var(\mathcal{F}_{P_{RM}}, +\infty)$ | $sign(\mathcal{F}_{P_{RM}}, +\infty)$ | Logically consistent? |
| --- | --- | --- | --- | --- | --- |
| 6 | $\{- + + - + - + -\}$ | N | 3 | ⋮ | ⋮ |
| 5 | $\{- + + - + - + +\}$ | N | 2 | $\{+ + - - + + + +\}$ | N |
| | $\{- + + - + + - +\}$ | Y | | $\{+ + - - - + + +\}$ | Y |
| | $\{- + + - + - - +\}$ | N | | $\{+ + - - - - + +\}$ | N |
| | $\{- + + - - + - +\}$ | Y | | $\{+ + - - - - - +\}$ | N |
| 4 | ⋮ | ⋮ | 1 | $\{+ + - - - - - -\}$ | N |

**Table 4:** Sturm sequence inequality sets for the RNA aptamer-based circuit in [20]. The sign of the first polynomial is fixed at *x* → 0. *var*(ℱ, 0) = 4 is not a logically consistent set.

| $var(\mathcal{F}, 0)$ | $sign(\mathcal{F}, 0)$ | Logically consistent? | $var(\mathcal{F}, +\infty)$ | $sign(\mathcal{F}, +\infty)$ | Logically consistent? |
| --- | --- | --- | --- | --- | --- |
| 4 | $\{- + - + -\}$ | N | 1 | ⋮ | ⋮ |
| 3 | $\{- - + - +\}$ | Y | 0 | $\{+ + + + +\}$ | Y |
| | $\{- + - + +\}$ | Y | | | |
| | $\{- + + - +\}$ | Y | | | |
| | $\{- + - - +\}$ | Y | | | |

We can compare the bistability region of the multi-operator P_RM_ system with that of a simple positive feedback circuit consisting of a dimeric repressor and only one operator site (MBN *kqw* in [13]). Rescaled as in Eq. (15), we have:

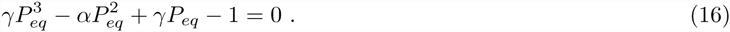

As with the MD toggle, this polynomial also has a maximum possible variation of 3 and thus only a single combination of inequalities that give rise to bistability, in the region given by

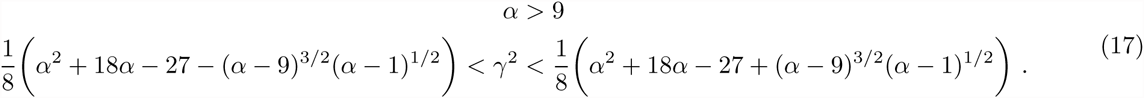

The bistable regions (in α–γ space) for these single gene circuits are shown in Fig. 2A. (The one-dimensional regions of bistability shown in bifurcation diagrams in [23] and [24] are plotted for comparison). It can be seen that the λ repressor circuit is bistable over a larger range and with lower values of the degradation rate constant. Interestingly, with α = 11 ([23] and references therein) the single operator circuit would just barely function as a bistable circuit, and any small fluctuation in circuit parameters would render it nonfunctional.

**Figure 2:**
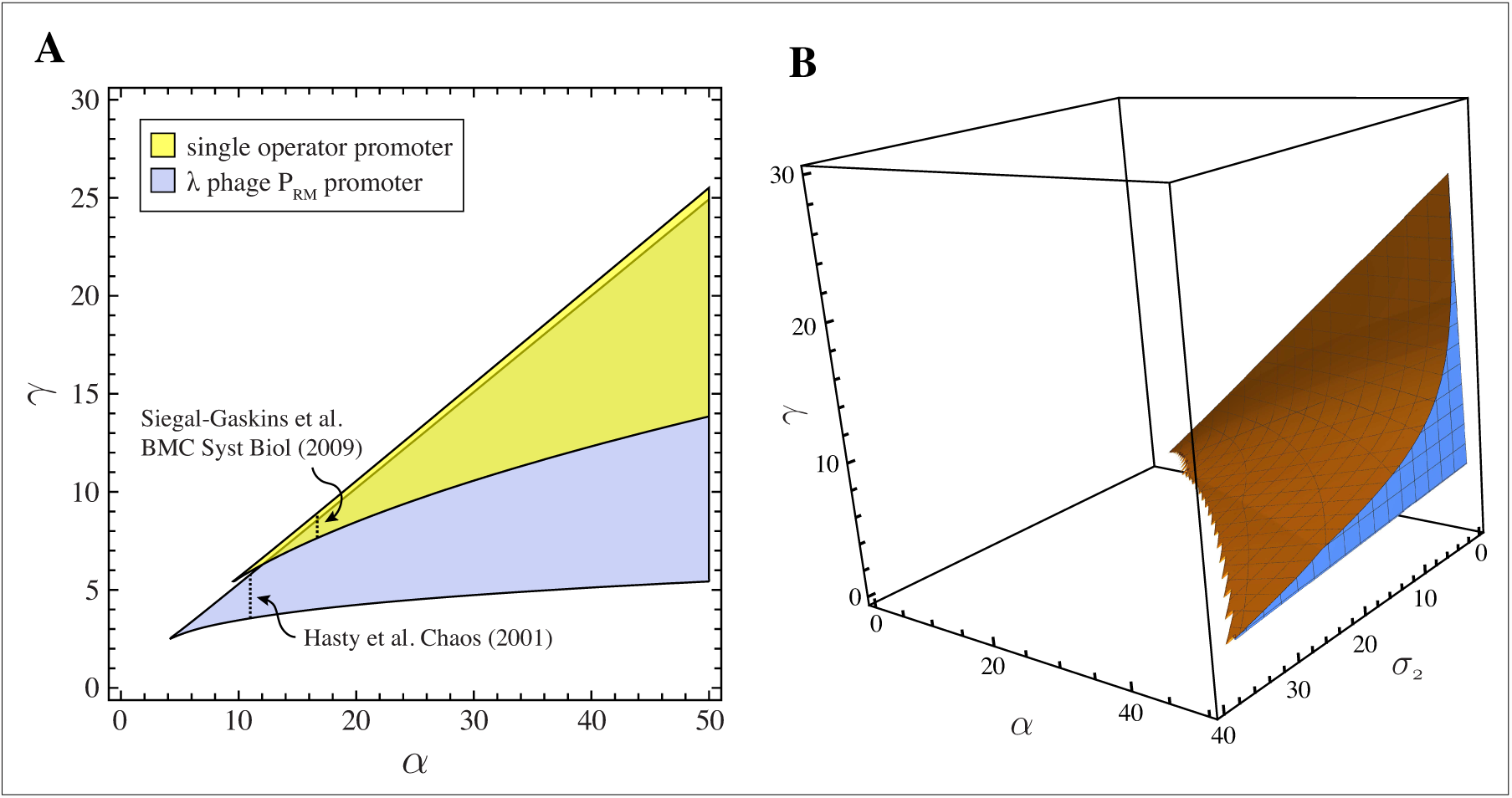
(A) Bistable regions for the single operator positive feedback circuit and the λ repressor–P_RM_ system with relative affinities σ_1_ = 2 and σ_2_ = 0.08. Vertical dashed lines indicate regions of bistability shown in the single-parameter bifurcation diagrams in [23] and [24]. (B) Region of bistability for the λ– P_RM_ system as a function of protein production enhancement α, protein degradation rate γ, and relative affinity for OR3 σ_2_.

We can also use our method to determine how the strength of the negative feedback (σ_2_) affects bistability. Keeping σ_1_ = 2, and using α = 11 and γ = 4.5 (centered in the bistable region at α = 11; see Fig. 2A), we find that σ_2_ can increase twelve-fold to ~ 0.96 before bistability is lost. In general, significant increases in σ_2_ require similar increases in α for bistability to be maintained, with the range of allowable α narrowing as a result (Fig. 2B).

## 4.3 Purely-monomeric toggle circuits

In a recent study [18] it was shown that a toggle variant consisting of two monomeric repressors derived from TALE protein DNA-binding domains cannot support bistability, but that the introduction of positive feedback in the form of two TALE-based activators that compete with the repressors for promoter access makes the system bistable. However, as we show below, it is not necessary to have two symmetric positive feedback loops for bistability; a single activator added to the monomeric toggle is sufficient.

We begin by combining components from the “mutual repressor” and “competitive feedback” models of [18] so that the two types of promoters—one containing a single binding site for both an activator and a repressor, and a second that is only responsive to a repressor—are present in one hybrid circuit:

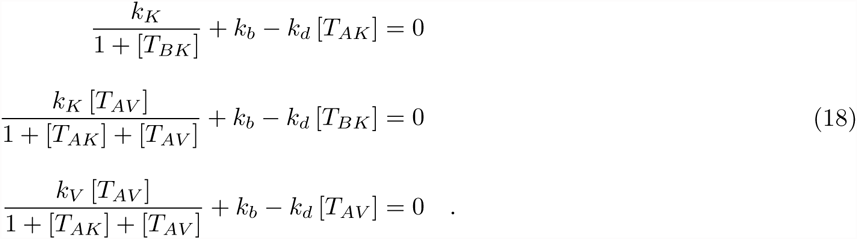

In keeping with the nomenclature of [18], [*T*_*AV*_] represents the concentration of the activator species, and [T_*AK*_] and [*T*_*BK*_] are the concentrations of the two repressor species. (For simplicity we have not included terms related to the fluorescent reporters as they have no effect on the capacity for bistability.) So that our results may be easily comparable to those in [18], we include a basal rate constant k_*b*_ to account for expression from the pristinamycin-and erythromycin-inducible promoters incorporated in the original system for external control. Other parameters of the models in [18] set to either zero or one in their simulations are, for clarity of presentation, also set to those same values here.

Eqs. (18) may be combined into a single equilibrium polynomial in [*T*_*AK*_]:

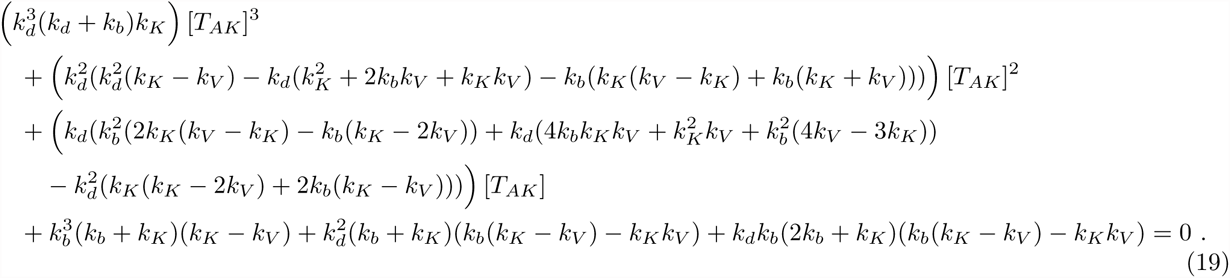

The maximum possible variation that the Sturm sequence associated with Eq. (19) may have is 3 and there is thus only a single combination of inequalities that can yield three steady states.

To establish the relationship between the repressor and activator expression strengths (k_*K*_ and k_*V*_, respectively) and the capacity for bistability, we set k_*b*_ to the uninduced steady-state value of ∼0.04 M·h^-1^ and the degradation rate constant k_*d*_ = 0.1 h^-1^ (values taken from [18]). Interestingly, we find two unconnected bistable subregions: one in which the repressor promoter strength must be greater than 10× that of the activator, and another in which the activator promoter strength must be greater than 2 that of the repressor (Fig. 3). Thus, while a purely monomeric toggle without positive feedback cannot be bistable—indeed, the equilibrium polynomial for such a circuit is only second order and could never have more than two real roots—the addition of a single activator with competitive feedback is all that is needed for bistability. Also noteworthy is that the values of k_*K*_ and k_*V*_ that give bistability in the simulation of the full “competitive feedback” model of [18] (180 h^-1^ and 30 h^-1^, respectively) only give a monostable response in this hybrid circuit.

**Figure 3:**
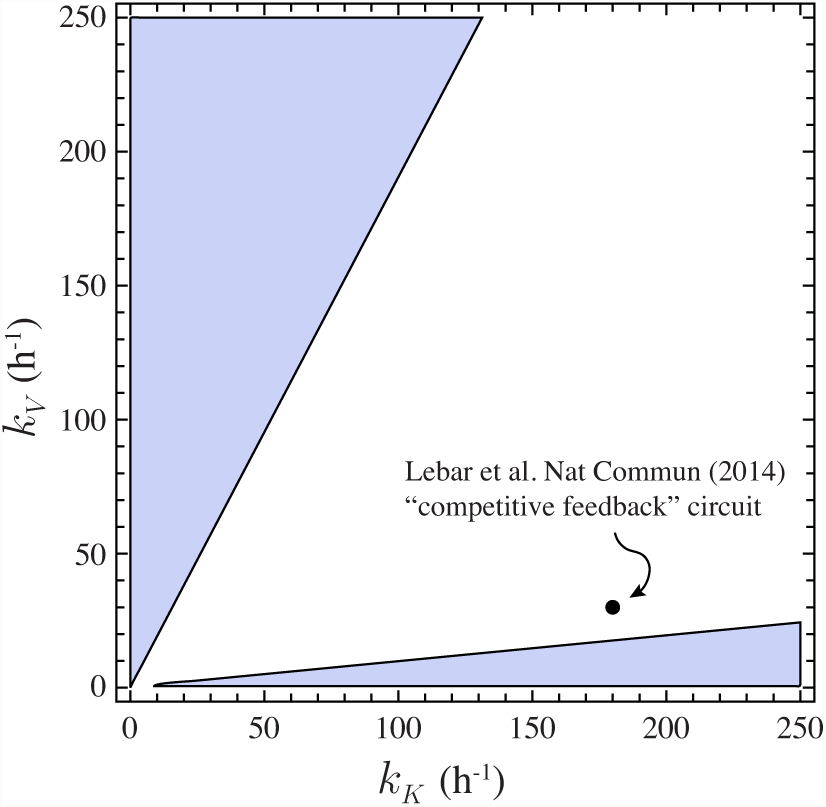
A monomeric toggle circuit with a single additional activator—a hybrid of the “mutual repressor” and “competitive feedback” models of [18]—yields two unconnected regions of bistability. The single point tested in the simulation of the full “competitive feedback” model in [18] is shown for comparison.

## 4.4 Sturm’s theorem and the Routh-Hurwitz stability criterion:a comparison of analytical methods

The Routh-Hurwitz stability criterion is commonly employed by control theorists to test if a polynomial admits positive roots. Typically applied to characteristic polynomials of linear time invariant systems to establish their stability, the method can also be used to find the parameter range in which a polynomial equilibrium condition admits a desired number of positive roots. Indeed, Sturm’s theorem and the Routh-Hurwitz method are related, and the validity of the latter can be demonstrated using Sturm chains. It may be argued that building a Routh table is a simpler procedure than applying Sturm’s theorem since it only requires basic arithmetic operations on the polynomial coefficients; however, its relative simplicity comes at a price: information about the real or complex nature of the positive roots is lost. In contrast, Sturm’s theorem allows for an exact determination of the number of real positive roots, as well as whether or not they are physically meaningful for a biochemical system.

A recent study of a novel biomolecular circuit that uses modulation of enzyme activity by inhibitory RNA aptamers to achieve bistability [20] provides us with an opportunity to compare our approach with one based on the Routh-Hurwitz. Interestingly, like the monomeric circuit from [18] and its asymmetric variant described above, and proposed MBN *bcdh* in [13], this aptamer-based circuit achieves bistability with monomeric regulation.

The equilibrium condition for the concentration of one of the enzyme species in its active form (given here by *x*; see Supporting Information) is a complicated polynomial of fourth order:

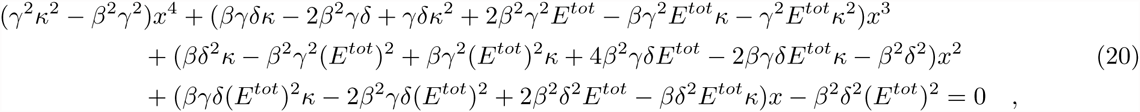

where E^*tot*^ is the total concentration of each of the two enzymes used in the circuit and Greek letters indicate reaction rates.

The inequality sets for this circuit are listed along with their allowabilities in Table 4.4. Combining these inequalities we find that for bistability it is necessary that

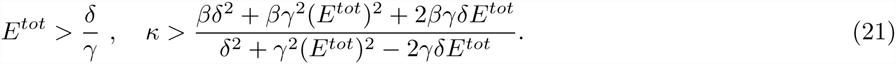

These conditions are identical to those identified in [20] by applying the Routh-Hurwitz table together with imposing conditions to guarantee *real* positive roots and conditions on the constant term of the characteristic polynomial to achieve bistability. Additional details are provided in the Supporting Information.

## 5 Discussion

In this work we have for the first time applied Sturm’s theorem to the study of biological circuits; specifically, biocircuits that can adopt two stable steady states. The primary results of our analyses are exact analytical expressions defining the various circuits’ regions of bistability, and without the need for parameter surveys and numerical simulation. We take advantage of the fact that the bistable circuits considered here are dissipative, positive in their linearization, and have characteristic polynomials with all non-constant terms positive, properties that allow us to ascertain the stability of their equilibria without computation. It is important to emphasize that these are not unique features of these particular circuits and that many real systems share these properties. And for any such system, once the number of steady-states is determined—by Sturm’s theorem or by other methods—the stabilities are known as well.

So following our analysis, can it be said that some topologies are in some sense “better” than others? Between the two different genetic toggle variants, we see that when rate parameters are fixed, the toggle consisting of two dimeric repressor species functions as a bistable switch over a wider range of DNA concentrations than one composed of one dimeric and one monomeric repressor. This result provides a strong motivation for choosing a DD toggle over a MD toggle in any application where there is considerable uncertainty or variability in parameter values or DNA concentrations (e.g., when DNA is in the form of plasmids without strict copy-number control). In single-gene positive feedback circuits exhibiting bistability, we see benefits to having additional operator sites: without both OR1 and OR2 in the λ repressor–P_RM_ system, the enhancement α = 11 would barely be sufficient for bistability. Interestingly, although feedback at OR3 is negative, it is not strong enough to significantly affect the circuit’s ability to function as a bistable switch. Taken together, these two results suggest that the promoter architecture of the λ system may have evolved to allow for both robust bistability due to the positive feedback as well as reduced variability or other benefit of the small negative autoregulation.

Though relatively little-known within the biological sciences, Sturm’s theorem has found applicability in a number of other areas where polynomials play an important role, including computational mathematics [25], dynamical systems [26, 27], robotics [28], and finance [29]. Additional biological applications are possible, for example as a tool to predict a large number of new bistable topologies or rule out those that do not have the capacity for bistability (like Chemical Reaction Network Theory, previously [13, 24]).

Other methods from algebraic geometry, including the closely related Descartes’ rule of signs, have also found use in biological systems analysis, with recent applications to model discrimination [30] and the study of chemical reaction networks [31, 32] and multisite phosphorylation systems [33], among many others. (This latter paper is particularly relevant to our own work, as it also highlights the benefits of treating parameters symbolically rather than numerically.) For the specific task required in this work—determining the number of positive real roots of a biocircuit’s equilbrium polynomial—Descartes’ rule of signs may be simpler to apply than Sturm’s theorem; however, it is definitively less powerful in that it can only give an upper bound on the number of real roots.

We note that not all bistable circuits of interest may be easily studied using our approach. Firstly, it may not be possible to capture the circuit’s equilibrium state with a single polynomial. When an equilibrium polynomial can be found, those of third-order are the simplest to study since there is only one combination of inequalities that can give rise to bistability. As the order increases, the number of inequality sets to test and the complexity of the Sturm sequences can also increase to the point where application of the theorem is impractical. However, this is highly dependent on the complexity of the polynomial coefficients and the number of Sturm sequences of fixed sign. Indeed, as we demonstrated with our analysis of the λ repressor–P_RM_ system, even a seventh-order polynomial can be tractable. For particularly large circuits for which an equilibrium polynomial cannot be derived, or for which the Sturm sequence is overly complex, a decomposition of the system into smaller modules can be beneficial.

The analysis of an equilbrium polynomial with a zero root—for example, that describing a recently-developed DNA-based circuit that achieves bistability through a combination of mutual inhibition and autocatalysis [34]—presents challenges, since Sturm’s theorem requires that neither of the limits of the region of interest (in our case, 0 and +∞) be roots. Under these circumstances one could choose an o < 0 for the lower limit; however, in this case there is no guarantee that the three roots found by Sturm’s theorem will all represent physically real, positive quantities (since one or more of the roots could in theory lie between o and 0).

## Extensions to Sturm’s theorem and related methods

The Sturm’s theorem approach described here can be directly applied to systems whose steady state may be characterized by a univariate polynomial with integer exponents and simple roots. While such systems are common, examples of equilibrium polynomials not of this particular form can also be found. For example, exponents that are typically given integer values in simplified models and/or to reflect strong multimerization (as in this work) or other reactions with significant positive cooperativity [40] can be found to have fractional values when the models are fit to empirical data [37–39]. In limited cases these generalised polynomials can be turned to a proper polynomial with a simple substitution that does affect the number of zeros (e.g., *u* = *x*^1^*/N*, if all exponents of x are multiples of 1/*N*). There are also simple extensions to Sturm’s theorem that deal with multiple roots (e.g., [35]) and multivariate models (e.g., [12, 36]). Should only an upper bound on the number of real roots of multivariate and generalised polynomial models be desired, extensions to the related Descartes’ rule of signs are also available [41–43].

## Acknowledgements

A large number of people contributed to this work with insights and comments. The authors would like to particularly thank Andras Gyorgy, Yutaka Hori, Scott C. Livingston, Anne Shiu, Eduardo Sontag, Elisenda Feliu, Zvi H. Rosen, Jaap Top, and Brian Ingalls. This research is funded in part by the National Science Foundation through Grant CMMI 1266402, and the Gordon and Betty Moore Foundation through Grant GBMF2809 to the Caltech Programmable Molecular Technology Initiative.

## S1 Equilibrium polynomial derivation for toggle variants and single-operator positive feedback circuit

The monomer-dimer toggle, dimer-dimer toggle, and single-operator positive feedback circuits were initially predicted to exhibit bistability using a chemical reaction network (CRN)-based topological survey [1]. Each of these CRNs contains reactions representing basal protein production and degradation:

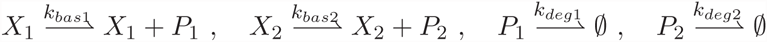

for genes *X*_*i*_ and proteins *P*_*i*_. The other reactions that uniquely define each circuit are:

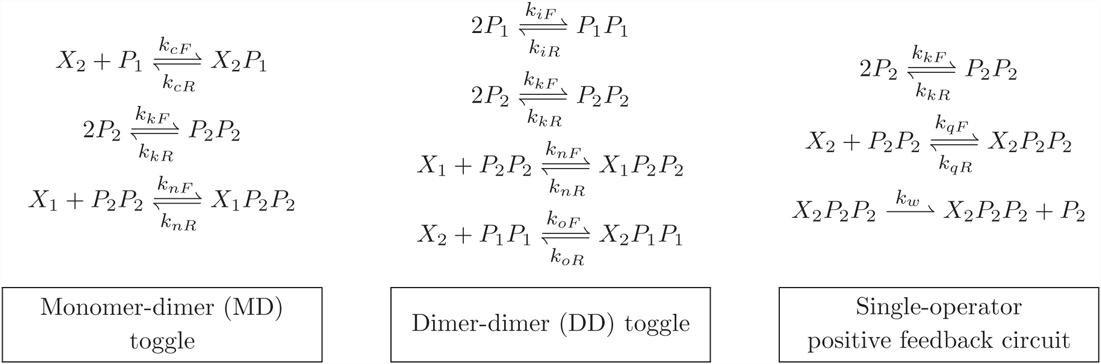

*P*_*i*_*P*_*i*_ represent dimeric species, and *X*_*i*_*P*_*j*_ and *X*_*i*_*P*_*j*_P_*j*_ represent monomers and dimers bound to the gene promoters. The various ODE sets were derived from these CRNs under the assumption of mass action kinetics and simplified using the fact that the total concentrations of each gene (in bound and unbound form) are conserved.

From these CRN formulations and assuming mass-action kinetics we can derive the following sets of ordinary differential equations (ODEs) that describe the circuit dynamics. For the MD toggle:

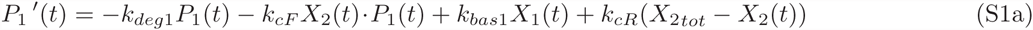

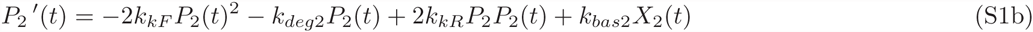

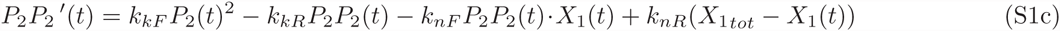

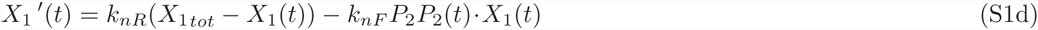

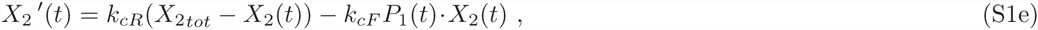

the DD toggle:

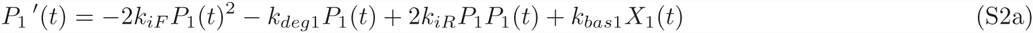

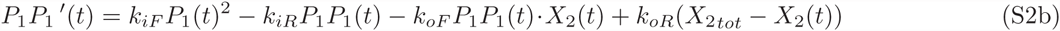

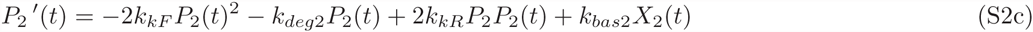

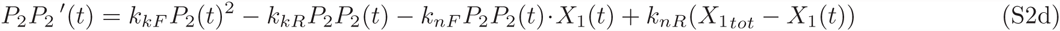

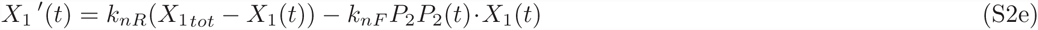

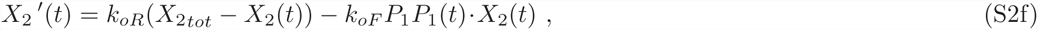

and single-operator positive feedback circuit:

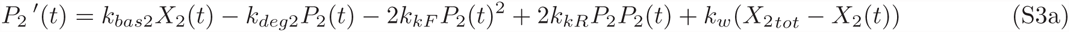

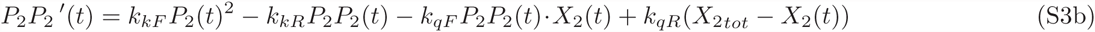

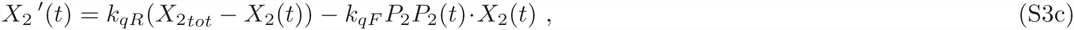

where the variable names are as in [1]: *X*_*i*_ is the concentration of free (i.e., unbound by repressor) gene i, *X*_*itot*_ is the total amount of *X*_*i*_ in the circuit (bound and unbound), and *P*_*i*_ and *P*_*i*_P_*i*_ represent the monomeric and dimeric forms of protein i, respectively. The various *k*_*x*_ are the reaction rates, and in the positive feedback circuit, *k*_*w*_ > *k*_*bas*2_ is assumed.

We derive the equilibrium polynomials from (S1), (S2), and (S3) by first setting the left-hand sides equal to zero. For the DD toggle, we subtract (S2d) from (S2e):

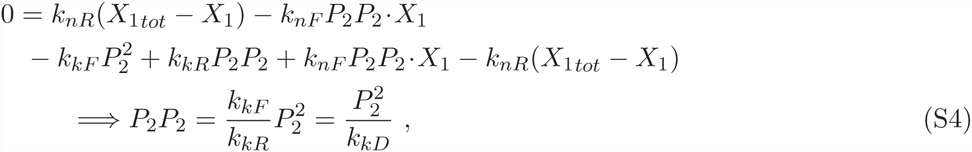

where for simplicity of notation we use *X*_*i*_, *P*_*i*_, and *P*_*i*_P_*i*_ to mean the equilibrium concentrations. We then plug this expression into (S2e) to get

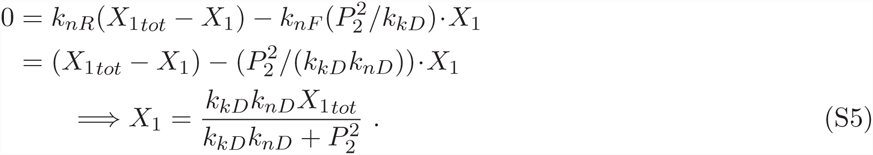

Similarly we subtract (S2b) from (S2f)

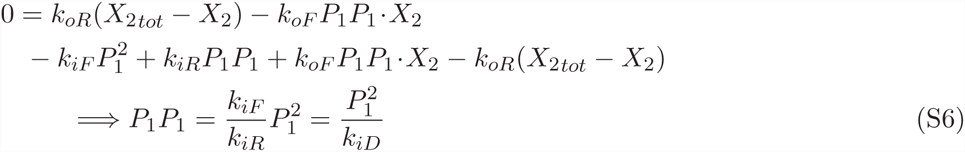

and plug the resulting expression back into (S2f) to get

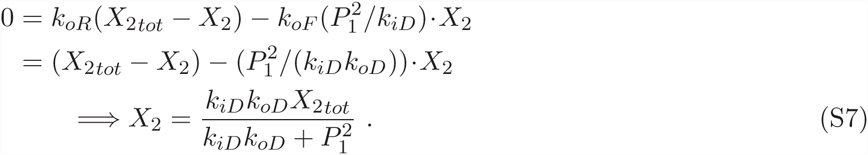

Substituting Eqs. (S4)–(S7) into (S2a) and (S2c) gives the equilibrium concentrations of *P*_1_ and *P*_2_ in the DD toggle shown in the main text:

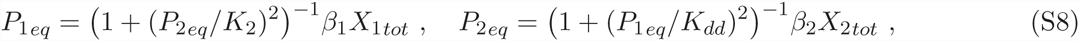

where *β_i_* = *k*_*basi*_/*k*_*degi*_ is the ratio of basal production and degradation rate constants for protein *i*, and the Michaelis constants *K*_*dd*_ = (*k*_*iD*_*k*_*oD*_)^1^*/*^2^ and *K*_2_ = (*k*_*kD*_*k*_*nD*_)^1^*/*^2^ represent the protein concentrations that yield 50% of the maximum production rate of their respective target proteins. The combination of these two expressions, with rescaling as described in the main text, gives the DD equilibrium polynomial.

For the MD toggle, we have again that

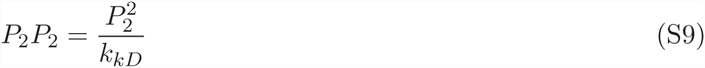

and

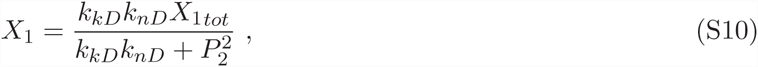

but also (from Eq. (S1e)) that

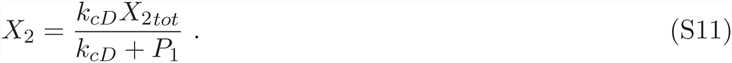

Substituting Eqs. (S9)–(S11) into (S1a) and (S1b) gives the equilibrium concentrations of *P*_1_ and *P*_2_ in the MD toggle shown in the main text:

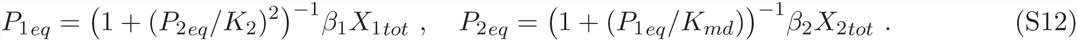

As previously, *β_i_* is the ratio of basal production and degradation rate constants for protein *i*, and the Michaelis constants *K*_*md*_ = *k*_*cD*_ and *K*_2_ = (*k*_*kD*_*k*_*nD*_)^1^*/*^2^ represent the protein concentrations that yield 50% of the maximum production rate of their respective targets. The combination of these two expressions, again with rescaling as described in the main text, gives the MD equilibrium polynomial.

The equilibrium polynomial for the single-operator positive feedback circuit may be similarly derived.

## S2 Aptamer-based bistable system with monomeric inhibition

The system described in [2] is composed by two enzymes *E*_1_ and *E*_2_ which transcribe at a constant rate two RNA aptamers *R*_1_ and *R*_2_. The aptamers mutually inhibit the enzymes; enzyme activity is recovered and aptamers are degraded with first order reaction rates. The reactions are:

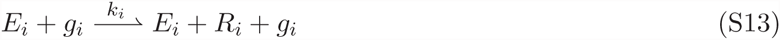

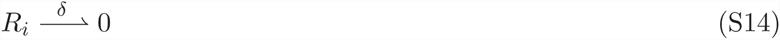

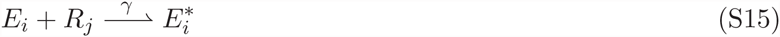

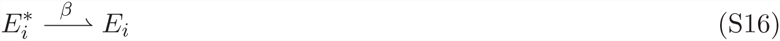

for i = 1,2 and j = 2,1, where *E*_*i*_ are active enzymes, 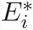 are inactive enzymes, *R*_*i*_ are RNA species, *g*_*i*_ are genes (constant). We assume the fraction of enzyme bound to its gene substrate is negligible. Using mass action laws, we can derive the ODEs describing the dynamics of the system:

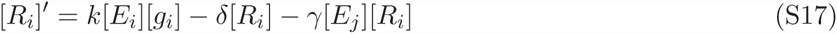

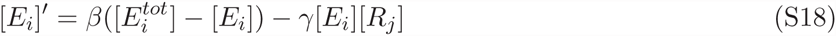

We switch to a more compact notation, and define *x*_1_ = [*R*_1_], *x*_2_ = [*E*_1_], *x*_3_ = [*R*_2_], and *x*_4_ = [*E*_2_]. Because the concentration of genes is constant, we define κ_1_ = *kg*_1_ and κ_2_ = *kg*_2_, and we assume κ_1_ = κ_2_ = κ. The system of ODEs becomes:

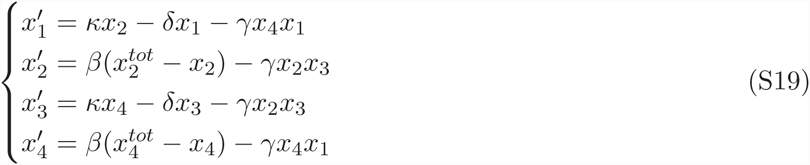

Equilibrium conditions are derived in detail in [2] from these expressions:

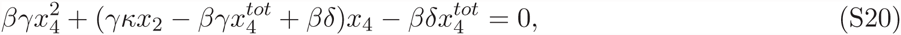

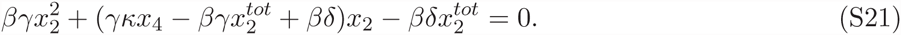

From here we can derive a fourth order polynomial in either *x*_2_ or *x*_4_:

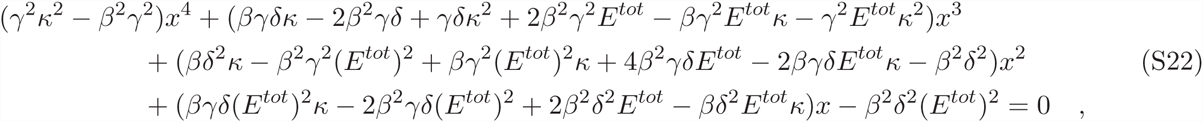

where we have used x to represent the steady-state concentration of an enzyme species in its active form and have put 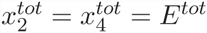 (as in [2]).

The inequality sets for this circuit along are listed along with their allowabilities in the main text (Table 6.4), as are the conditions on the parameters that yield bistability.

## S3 Sturm polynomials

Sturm polynomials can be rather long and complicated functions; however, they can be easily generated using software capable of symbolic manipulation (e.g., Mathematica). We present here the Sturm sequences (in x, for simplicity of notation) for the MD toggle, DD toggle, and single-operator positive feedback circuits.

The Sturm polynomials associated with the MD toggle equilibrium polynomial are:

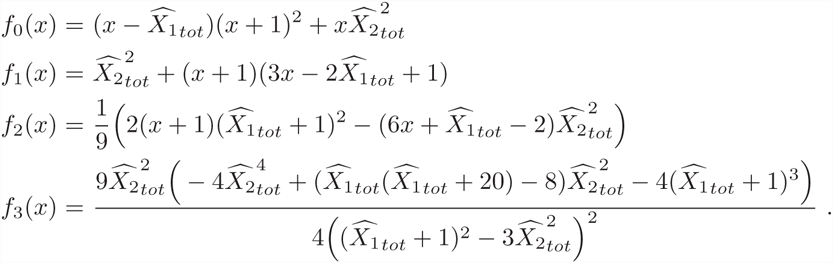

The DD Sturm polynomials are:

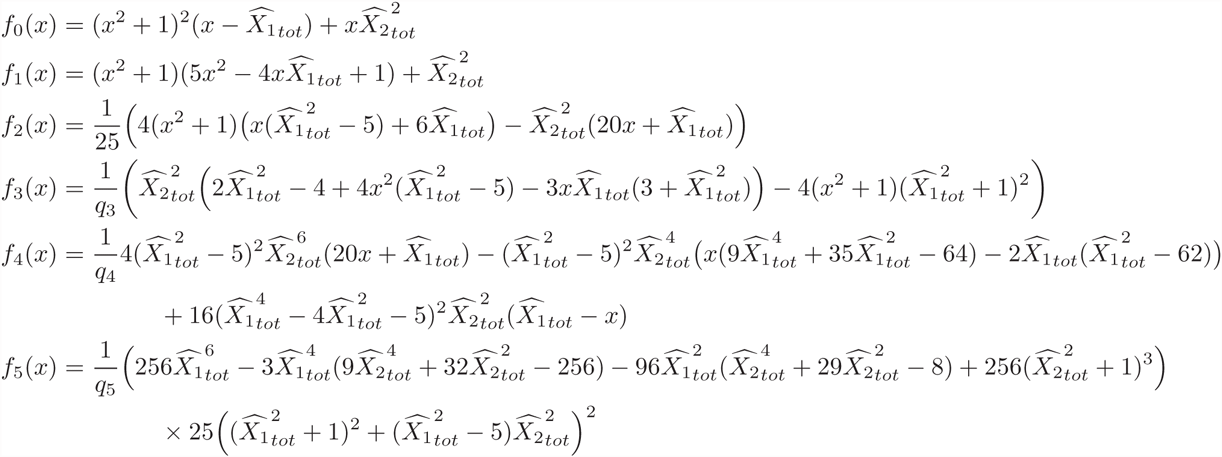

where

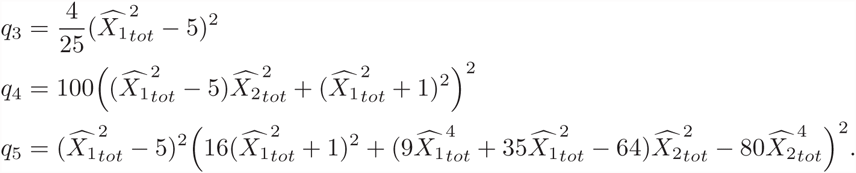

It can be seen that some of the polynomials in the DD toggle sequence contain a 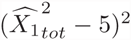 term which renders them indeterminate when 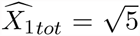. (The same term is responsible for polynomial *f*_4_(*x*) → 0 when *x* → 0, thus terminating the sequence prematurely.) To avoid these problems it is necessary to generate two different sequences for this toggle: one where we assume 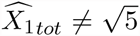 in Eq. (8), and another with 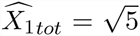. In this latter case the DD toggle equilibrium polynomial Eq. (8) becomes

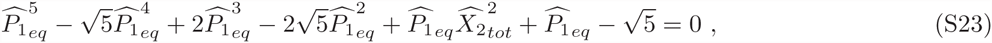

which we use to generate a second Sturm sequence for use at 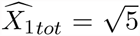 only:

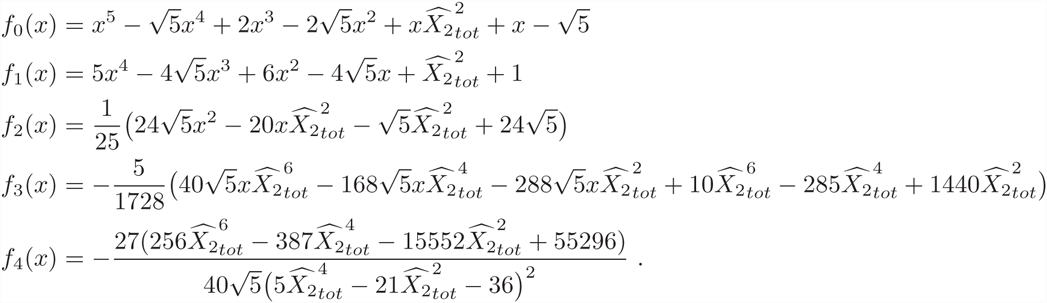

The single-operator positive feedback circuit Sturm sequence contains the following polynomials:

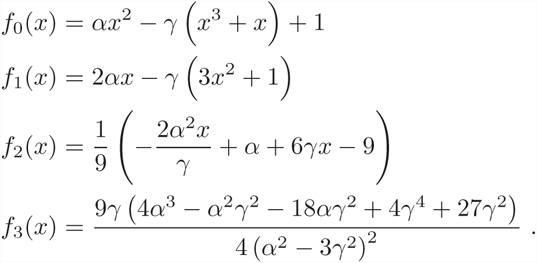

Recall that γ is the rescaled degradation rate constant for the λ repressor and α represents the increase in protein production resulting from repressor dimer binding to OR2.

## S4 Number of steady states and stability analysis

### 1 S4.1 Preliminaries

**Definition 1 (Positive systems)** *A linear system* 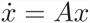 *is positive if for every nonegative initial state the solution* x(t) *is nonnegative.*

The following is a well known condition for positivity [3]:

**Theorem 1** *A linear system* 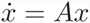 *is positive if and only if matrix* A *is a Metzler matrix, i.e., its elements satisfy:* a_*ij*_ ≥ 0, ∀(i, j) *such that* i ≠ j.

The Jacobian matrices of all systems considered here are, for any choice of parameters, similar to Metzler matrices via linear transformations. For example, the Jacobians J_*i*_ of each of the MD toggle, DD toggle, single-operator positive feedback, and aptamer-based inhibition circuits may be transformed to Metzler matrices 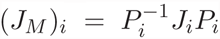, with *P*_*md*_ = diag(−1, 1, 1, −1, 1), *P*_dd_ = diag(−1, −1, 1, 1, −1, 1), *P*_*pf*_ = diag(1, 1, −1), and *P*_*apt*_ = diag(1, 1, −1, −1) respectively. Thus, the linearizations of the circuit ODEs are positive.

The general definition of *dissipativity* (see, e.g., [4]) is based on the existence of compact, forward invariant subsets of 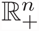 that absorb the system trajectories. The following definition (from [5]) is equivalent and easier to verify:

**Definition 2 (Dissipative systems)** *A system* 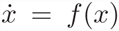 *is dissipative if its solutions are eventually uniformly bounded, i.e., there exists a constant k* > 0 *such that:*

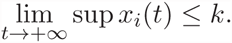

The systems analyzed in this work are all dissipative. As an example, we verify the definition for the MD toggle model (S1). Because the total mass of each of the DNA species *X*_1_ and *X*_2_ is constant, we know that *X*_1_(*t*) ≤ *X*_*max*_ and *X*_2_(*t*) ≤ *X*_*max*_ ∀ *t*, where *X*_*max*_ = max{*X*_1*tot*_, *X*_2*tot*_}. The concentration of *P*_1_ can be upper bounded as follows:

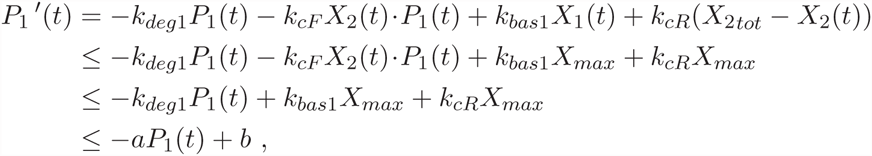

where *a* = *k*_*deg1*_ and *b* = (*k*_*bas*_1 + *k*_*cR*_)*X*_*max*_. The right hand side of the last inequality above is a linear, asymptotically stable system whose solution is eventually uniformly bounded (*b* is a finite constant). Using the comparison principle [6], we conclude that *P*_1_(*t*) is bounded and can find a constant *k* that satisfies the definition.

*P*_2_ may be similarly upper bounded. We first consider the dynamics of 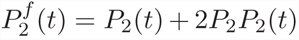, the total amount of unbound *P*_2_ in the system:

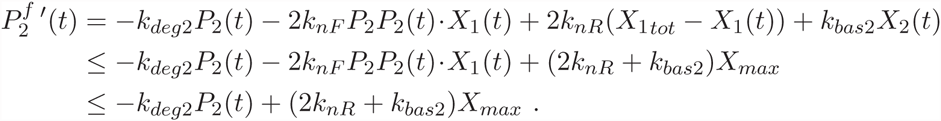

The dynamics of monomeric *P*_2_ satisfy:

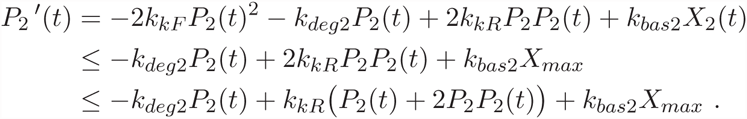

Together, we have:

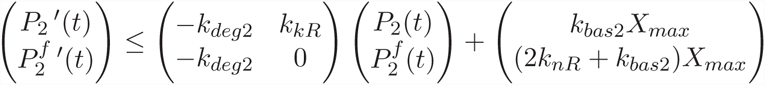

The variables 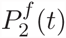 and *P*_2_(*t*) are upper bounded by a linear system with eigenvalues

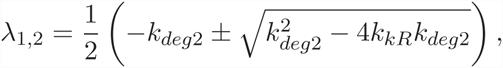

whose real part is always negative for any value of the (positive) parameters. Being upper bounded by an asymptotically stable linear system, the concentrations *P*_2_ and 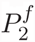 are eventually uniformly bounded. It follows that *P*_2_*P*_2_ is also eventually uniformly bounded, since 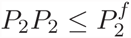.

Therefore, the ODE model of the monomer-dimer toggle system is dissipative. Note that the same conclusion cannot be reached in the absence of degradation (*k*_*deg*2_ = 0) since the total amount of protein will grow unbounded.

We can also easily show that system (S19) is dissipative: the total concentration of enzymes is constant, thus bounded at all times. The RNA species are also asymptotically bounded. The dynamics of RNA *x*_1_, for instance, can be upper bounded by an asymptotically stable linear time invariant system:

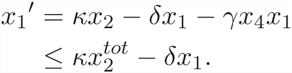

We can derive a similar upper bound for 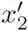.

The other systems considered in this work can be similarly shown to be dissipative.

### S4.2 Stability of equilibria

Sturm’s theorem applied to our circuits’ polynomial equilibrium conditions reveals that each system admits three positive equilibria. The stability properties of these equilibria can be determined by *degree theory* [5].

**Definition 3 (Regular equilibrium)** *An equilibrium point 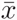 of system 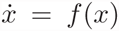 is regular if* 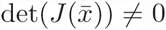 *(in other words, 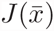 must be invertible; alternatively, 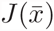 must not have eigenvalues at the origin).*

Definition 4 (Index of an equilibrium point) *The index of a regular equilibrium point 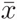 is the sign of the determinant of* 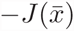:

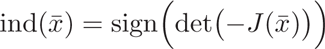

**Definition 5 (Degree of a system)** *The degree of a dynamical system* 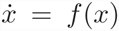 *having equilibria* 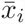, *i* = 1, …, *m, is defined as:*

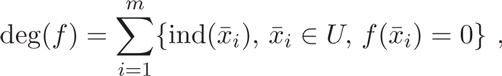

*where 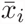 are regular equilibria.*

**Theorem 2** *A dissipative dynamical system 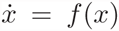 defined on* ℝ^*n*^ *has degree +1 with respect to any bounded open set containing all its equilibria.*

Since the systems are all dissipative, by Theorem 2 all have degree +1. We further note that the Jacobian matrices are row equivalent to the identity matrix (as confirmed with Mathematica) and thus always invertible—det(*J*) ≠ 0 for all (positive) parameters and equilibria. Therefore, all equilibria are regular. To determine the index of each equilibrium point, we need not know the value of the equilibrium itself, since in general

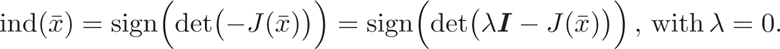

Therefore, the index of an equilibrium corresponds to the sign of the constant term in the system’s characteristic polynomial 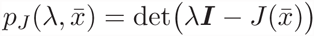.

For the Jacobians under consideration, the sign of the constant term in the characteristic polynomial also determines the stability properties of the corresponding equilibrium; we can state the following lemma [2]:

**Lemma 1** *Any single equilibrium 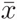 of a system under consideration in this work is unstable if and only if the constant term of the characteristic polynomial* p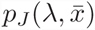) *is negative. Instability can only be driven by a simple, real (positive) eigenvalue.*

**Proof** The linearizations of our circuit ODEs define positive linear systems, where the Jacobians are similar to Metzler matrices. Therefore, these Jacobians always have a real dominant eigenvalue, i.e. λ_*max*_ ≥ ℛ*e*(λ_*i*_), ∀λ_*i*_ ∈ J [7].

The coefficients of the characteristic polynomials p_*J*_ (λ, 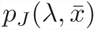) are all real and positive with the exception of the constant terms, which can be positive or negative. If a constant term is negative, then we know that 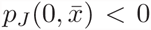 and it is real. In the limit λ → ∞, 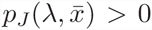 > 0 because all other coefficients are positive. Thus, there must be at least one point in the right half plane that is a root of 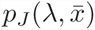, the system is unstable, and because the Jacobian is similar to a Metzler matrix, the largest root must be real.

If a system is unstable, then its characteristic polynomial must have at least one root with positive real part. *Ab absurdo*, suppose the constant term is positive. Then instability can only occur with a pair of complex conjugate eigenvalues with positive real part. This is impossible because the Jacobian is a Metzler matrix and the dominant eigenvalue must be real. Thus, the constant term of the characteristic polynomial must be negative. ▪

We can now finish our stability analysis. Our systems all have degree +1 (Theorem 2), thus when three equilibria are present their indices must be equal to +1, +1, and -1 so that their sum is +1 (we recall that all equilibria of our systems are regular). Since the index is equal to the sign of the constant term in the characteristic polynomial, a positive index is associated with a stable equilibrium and a negative index is associated with an unstable equilibrium, and we can conclude that, with three equilibria, our systems are bistable. Note that the unstable point does not admit local oscillatory behaviors, because local instability is driven by a real eigenvalue (Lemma 1). As an alternative argument, we can also simply note that our systems are monotone—for any choice of parameters the Jacobians are similar to Metzler matrices, a property that defines a monotone system with respect to the positive orthant [8, 9]—and a monotone system does not admit oscillatory behaviors.

